# Converging PMF calculations of antibiotic permeation across an outer membrane porin with sub-kilocalorie per mole accuracy

**DOI:** 10.1101/2023.03.27.534415

**Authors:** Jeremy Lapierre, Jochen S. Hub

## Abstract

The emergence of multi-drug resistant pathogens led to a critical need for new antibiotics. A key property of effective antibiotics against Gram-negative bacteria is their ability to permeate through the bacterial outer membrane via transmembrane porin proteins. Molecular dynamics (MD) simulations are in principle capable of modeling antibiotic permeation across outer membrane porins (OMPs). However, owing to sampling problems, it has remained challenging to obtain converged potentials of mean force (PMFs) for antibiotic permeation across OMPs. Here, we investigated the convergence of PMFs obtained with three advanced flavors of the umbrella sampling (US) technique aimed to quantify the permeation of the antibiotic fosmidomycin across the OprO porin: (i) Hamiltonian replica-exchange with solute tempering in combination with US, (ii) simulated tempering-enhanced US, and (iii) replica-exchange US. To quantify the PMF convergence and to reveal hysteresis problems, we computed several independent sets of US simulations started from pulling simulations in outward and inward permeation directions. We find that replica-exchange US in combination with well-chosen restraints is highly successful for obtaining converged PMFs of fosmidomycin permeation through OprO, reaching PMFs converged to sub-kilocalorie per mole accuracy.

**TOC Graphic:** 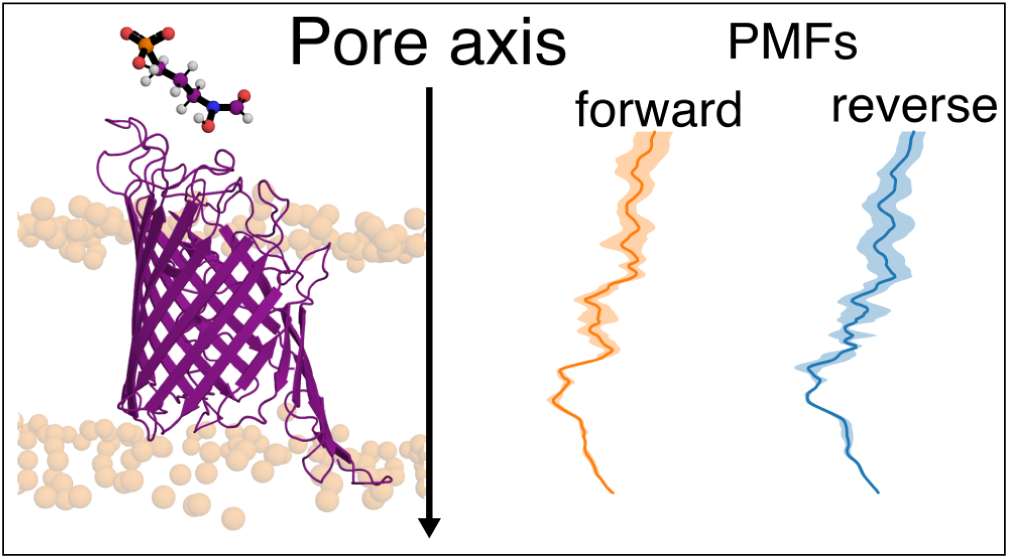

## Introduction

*Pseudomonas aeruginosa* are Gram-negative bacteria, which differ from Gram-positive bacteria by the presence of an outer membrane and of a thinner peptidoglycan layer (Fig. 1). *Pseudomonas aeruginosa* is an opportunistic pathogen, implying that it is usually not harmful to healthy individuals but may cause severe disease in hosts who suffer from a defective immune system or who are weakened by another diseases. For this reason, infections by *Pseudomonas aeruginosa* are frequent in hospitals and, accordingly, classified as nosocomial infections.^1–3^ Pathogens causing nosocomial infections are often multi-drug resistant, this is specifically true for *Pseudomonas aeruginosa*. The World Health Organization emphasized a critical need of new antibiotics against this group of pathogens to better protect hospitalized patients.^4^

**Figure 1:**
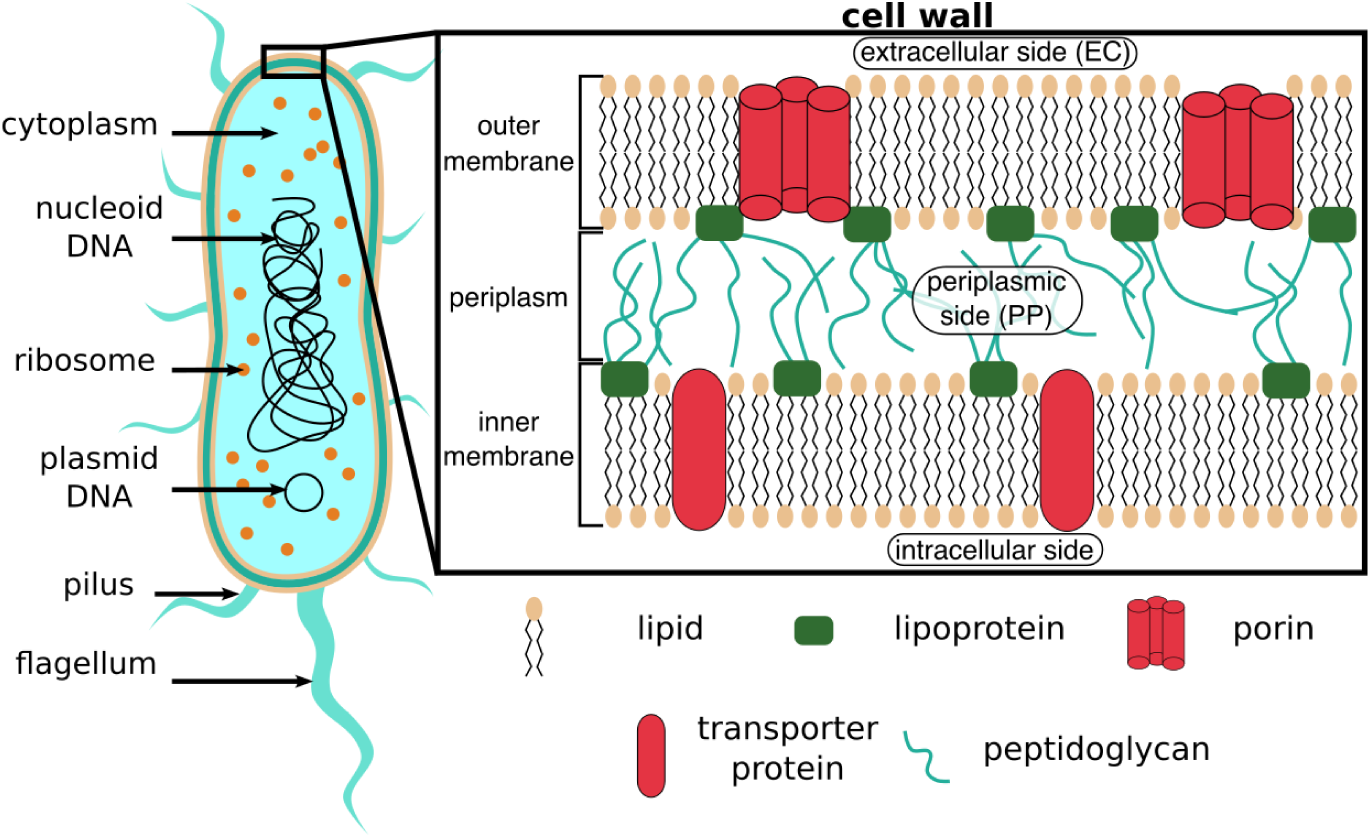
Overall structure and macromolecular components of *Pseudomonas aeruginosa*, a gram-negative bacterium. Outer membrane porins (red cylinders) constitute possible uptake pathways for antibiotics. {fig:bacteria}

Absorption of nutrients by *Pseudomonas aeruginosa* from the extracellular medium is mediated by outer membrane porin proteins (Fig. 1), which select incoming molecules by their charge and size.^5–7^ To design new effective antibiotics, it is thus of paramount importance to take into account the permeability porins for drug canidates. Notably, *Pseudomonas aeruginosa* lacks many porins found in other Gram-negative bacteria, which has been associated with particular drug-resistance, ^7–10^ although recent work emphasized the importance of direct permeation across the lipid membrane.^11^ Detailed descriptions of bacterial outer membrane porins as well as permeation mechanisms of small-molecules through these molecular gateways have been reviewed in detail.^12, 13^

In this study, we focused on the permeation of fosmidomycin through the OprO porin, a polyphosphate-specific homotrimeric transmembrane protein. Fosmidomycin is an inhibitor of the 1-deoxy-D-xylulose 5-phosphate reductoisomerase, an enzyme involved in a biosynthesis pathway for isoprenoids, but specific to bacteria and protozoa. Isoprenoids play major roles in functions such as electron transport or cell signaling,^14^ therefore blocking the bacterial specific pathway of isoprenoid biosynthesis is an effective strategy to impede proliferation of the pathogen.

Atomistic molecular dynamics (MD) simulations have been used to study drug permeation across biological membranes, however, such simulations are subject to considerable sampling challenges. Golla *et al.* recently showed that potential of mean forces (PMF) calculations based on standalone umbrella sampling (US)^15^ using the drug position along the pore as collective variable (CV) suffers from major hysteresis effects,^16^ whereas well-tempered metadynamics with multiple walkers along the same CV yielded more accurate PMFs.^16, 17^ In line with the work from Golla *et al.*,^16^ well-tempered metadynamics with multiple walkers have also been successfully applied for studying permeation processes of several small molecules through the OmpF porin from *E. coli* .^18^ Because standalone US frequently suffers from sampling and hysteresis problems, several improved flavors of US have been developed including Hamiltonian replica-exchange US, ^19–21^ temperature-accelerated sliced sampling (TASS),^22^ and simulated tempering-enhanced umbrella sampling (STeUS).^23^ In fact, replica-exchange US —a specific application of Hamiltonian replica-exchange simulations— has been successfully applied to obtain quantitative insight into antibiotic permeation through OmpF.^24, 25^ Acharya *et al.* applied TASS to rationalize permeation of ciprofloxacin through OmpF.^26^ Furthermore, Vasan *et al.* analyzed the role of large-scale loop transitions in OmpF, which pose particular sampling challenges. ^25^ Despite these recent achievements, obtaining converged PMFs of antibiotic permeation across porins remains a challenge. Furthermore, since systematic comparisons of augmented US variants are still limited in the literature, finding an optimal US protocol for computing converged PMFs for drug permeation is time-consuming.

In this work, we have compared four methods for computing PMFs of the permeation of fosmidomycin through the OprO porin: (i) standalone US,^15^ (ii) US augmented with Hamiltonian replica-exchange with charge-scaling, and more specifically with the related REST2 method (US-HREX),^20, 21^ (iii) simulated tempering-enhanced umbrella sampling (STeUS),^23, 27^ and (iv) replica-exchange umbrella sampling (REUS, also called bias-exchange US, BEUS).^19^ Whereas OprO forms trimers under native conditions (Fig. 1, red cylinders), we here simulated only the monomer to facilitate the computational setups and justified by the fact that we focused finding an efficient sampling method rather than reproducing the native permeation process (Fig. 2A).

We identified four key methodological ingredients that were critical for obtaining converged PMFs of the permeation of fosmidomycin through OprO: (i) the use us REUS to enhance sampling during US simulations; (ii) the application of orientational flat-bottomed restraints to maintain one pre-selected orientation of the elongated solute inside the channel; (iii) the use of cylindrical flat-bottomed restraints to keep the solute in the proximity of the pore axis; and (iv), during the initial constant-velocity pulling of the solute along the channel, the application of position restraints on protein atoms to avoid distortions of pore-lining residues. By combining these ingredients, we obtained PMFs that were converged with an accuracy below one kilocalorie per mole.

## Methods

### Simulation setup

REUS and US-HREX were carried out with Gromacs^28^ version2020.6 built with Open MPI and patched with Plumed version 2.7.2.^29, 30^ Standalone US and STeUS were carried out with Gromacs version 2020.4 patched with Plumed 2.7.0. Atomic coordinates of the porin trimers (pdb code 4RJW^31^) and fosmidomycin, as well as forcefield parameters were kindly provided by Prof. Ulrich Kleinekathöfer.^16, 17^ Protein, lipids, water and ions were parameterized with the CHARMM36m forcefield^32^ and fosmidomycin with the CGenFF-based forcefield generated with the ParamChem server.^33^ The validation of fosmidomycin parameters are available in Ref. 16. From atomic coordinates of OprO trimers, two monomers have been removed to obtain a monomeric form of the OprO porin. We inserted the monomer into a lipid bilayer of 334 1-palmitoyl-2-oleoyl-*sn*-glycero-3-phosphoethanolamine (POPE) lipids with the CHARMM-GUI membrane builder.^34^ We solvated the system with TIP3P water molecules,^35^ and added 14 potassium ions to neutralize the system. For simplicity, the effect of additional salt was not considered in this study.

Electrostatic interactions were computed with the particle-mesh Ewald method, ^36^ using a Fourier grid spacing of 0.12 nm and a real-space cutoff at 1.12 nm. Short-range repulsion and dispersion interactions were described with a Lennard-Jones potential with a cutoff at 1.2 nm and a force-switch modifier set to 1.0 nm. Angles and bonds of water molecules were constrained with SETTLE,^37^ and bonds involving other hydrogen atoms were constrained with LINCS.^38^ Energy minimization was carried out with steepest descent and equilibration was performed following the six-step protocol provided by CHARMM-GUI.^34^ In brief, the aforementioned equilibration protocol consisted in two 125 ps NVT equilibration simulations, one 125 ps NPT equilibration, and three 500 ps NPT simulations. During equilibration, position restraints were applied on lipid phosphate atoms, and to protein and fosmidomycin heavy atoms; restraints were slowly released through the six equilibration steps. Pressure at 1 bar was controlled by the Berendsen barostat (*τ* =5 ps) and temperature at 300 K by the Berendsen thermostat (*τ* =1 ps).^39^

For production runs, a 4 fs time step was used for standalone US, US-HREX and REUS, and a 3.5 fs time step was used for STeUS. Using a time-step larger than the commonly used 2 fs time step was possible by modelling all hydrogen atoms as virtual sites. The pressure was controlled with the Parrinello-Rahman barostat,^40^ while the temperature was controlled by velocity rescaling. ^41^

### Standalone umbrella sampling

To obtain starting conformations of US, we carried out eight 100 ns independent constant-velocity pulling simulations (force constant 1000 kJ mol*^−^*^1^ nm*^−^*^2^). We pulled fosmidomycin from the extracellular side (EC) to the periplasmic side (PP) (forward direction) or from PP to EC (reverse direction), each direction with two fosmidomycin orientations as defined in Fig. 2C, and simulating two repetitions for each setup (2 × 2 × 2 pulling simulations in total). During pulling simulations, we pulled along the *z*-component of the vector connecting (i) the center of mass (COM) of C*_α_* atoms close to the cavity and (ii) the COM of the terminal chemical moiety of fosmidomycin in the direction of the movement. For the terminal chemical moieties, we used either phosphoryl group (PP-to-EC in orientation 1; EC-to-PP in orientation 2) or the amine group (PP-to-EC in orientation 2; EC-to-PP in orientation 1). During pulling simulations, we used three types of restraints, which were critical to obtain converged PMFs, as described below. Trajectories from pulling simulations were post-processed to map each frame onto the CV *z* defined as the *z*-component of the vector connecting the COM of OprO porin C*_α_* atoms close to the cavity, and the center of mass of the entire fosmidomycin.

**Figure 2:**
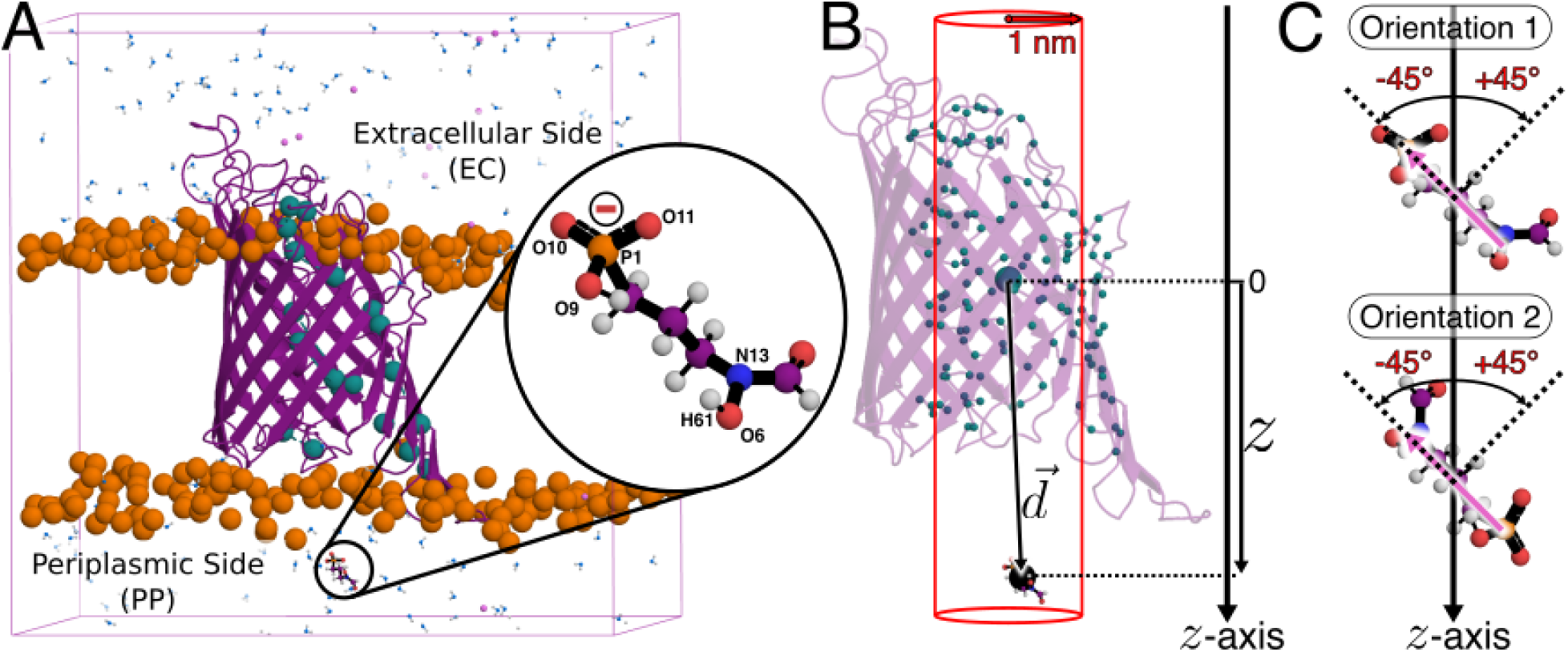
Setup for simulating fosmidomycin permeation through the OprO porin. (A) Simulation system. OprO is shown as purple cartoon, POPE phosphate groups as orange spheres, key lysine and arginine residues along the porin contributing to its anionic selectivity are highlighted with cyan spheres, fosmidomycin and few water molecules are represented as balls and sticks, and potassium cations are shown as small pink spheres. Most of water molecules have been removed for clarity. The focus on fosmidomycin shows key atoms that have been used for defining the CV. (B) The large cyan bead depicts the center of mass of all C*_α_* atoms close to the porin’s lumen (pCOM) represented as small cyan beads. The large black sphere represents the center of mass of the following atoms in fosmidomycin: P1, O9– 11, N13, H61 and O6 (fCOM). The *z*-component of the vector connecting the pCOM and the fCOM was used as CV for driving the permeation process. The red cylinder illustrates the cylindrical flat-bottomed position restraint. (C) Two fosmidomycin’s orientations studied in this work. The pink arrow represents the vector connecting COMs of phosphoryl and amine group; the angle between this vector and the *z*-axis has been restrained with a flat-bottom potential such that this angle is maintained within [−45*^◦^,* +45*^◦^*]. {oproview}

After mapping frames onto *z*, we launched 144 umbrella simulations in the range *z* ∈ [−3.4, 3.34] nm, with a force constant of 2000 kJ mol*^−^*^1^ nm*^−^*^2^ for 30 ns. During this equilibration step, force constants of the restraints on the backbone and side chains heavy atoms of the protein were reduce to 100 kJ mol*^−^*^1^ nm*^−^*^2^ and 10 kJ mol*^−^*^1^ nm*^−^*^2^ respectively. Final coordinates obtained from the previous step were then used to launch production simulations of 200 ns. During production, position restraints on protein backbone and protein side chains were completely removed. Centers of harmonic potentials used in all US flavors are provided in Fig. S1. The choice of umbrella window spacing is described below in the section on REUS.

All PMFs were computed with wham by Alan Grossfield (“WHAM: the weighted histogram analysis method”, version 2.0.11, http://membrane.urmc.rochester.edu/wordpress/?page_id=126).

### Combining umbrella sampling with Hamiltonian-replica exchange: US-HREX

The CV, number of umbrella windows, spacing between windows, initial configurations, restraints used during production, and US force constant were chosen as during standalone US. To enhance the sampling in higher replicas (corresponding to lower *λ* values), we scaled charges as depicted in Fig. S2A. The protocol used to scale positive and negative charges is shown in Fig. S2B.

To identify the *λ*-range that maintains a stable protein conformation, we ran six simulations for 400 ns with the following *λ* values: 1, 0.8, 0.6, 0.4, 0.2, 0.1 and 0.05. Simulations with *λ* = 0.1 or 0.05 were numerically unstable. However, simulations with *λ* = 1 through 0.2 were stable over 400 ns, and visual inspection of trajectories as well as root mean square residue fluctuations did not indicate protein unfolding. For production, we ran 24 *λ*-replicas for each of our 144 umbrella windows. In order to obtain ∼ 19% of exchange acceptance between the 24 replicas, we scaled positive charges from *λ* = 1 to *λ* = 0.793 in steps of 0.009. Each umbrella window was simulated for 9 ns, leading with 24 replicas to an overall simulation time of 216 ns per window. Thus, the total simulation time per US window was similar to the simulation time of 200 ns used for other US flavors.

### Combining umbrella sampling with simulated tempering: STeUS

The CV *z*, number of umbrella windows, initial configurations, restraints used during production, total simulation time, and US force constant were chosen as during standalone US. Simulated tempering was carried out with temperatures ranging from 300 K to 348 K in steps of 4 K. We chose the initial temperature weights following Park *et al.*,^42^ which involves simulated annealing simulation from the lowest to the highest temperature. Accordingly, the weights were chosen as

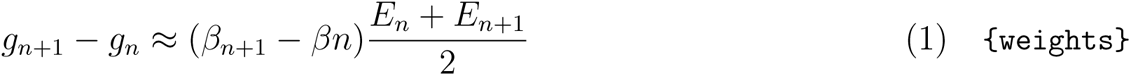

where *g_n_* is the weight of the state with temperature *T_n_*, *β_n_* = 1*/k_B_T_n_* is the respective inverse temperature, and *E_n_*is the average potential energy for temperature state *T_n_*. To obtain the *E_n_*, we carried out a simulated annealing simulation for the umbrella window restrained to *z* = 0 nm, for which the temperature was increased from 300 K to 348 K, with 2 ns per temperature state and 100 ps for each heating step over a total simulation time of 27.2 ns. With this protocol, we obtained the following weights from *g*_12_ through *g*_0_: 0, 4370.5, 8600.3, 12694.8, 16659.9, 20500.2, 24220.5, 27826.5, 31322.1, 34711.3, 37999.4, 41189.1, 44284.0.

To further optimize the weights, we carried out a simulated tempering simulation for umbrella window corresponding to *z* = 0 nm with the previously determined weights, with an exchange attempt every 100 steps and using the Wang-Landau algorithm over a total simulation time of 43 ns. This simulation was used to determine the final weights used for production: 0, 4361.5, 8577.3, 12657.8, 16617.9, 20459.2, 24166.5, 27768.5, 31265.1, 34644.3, 37926.4, 41114.1, 44206.0. Finally, each of the 144 umbrella windows has been run with simulated tempering using final weights determined as explained above, with exchange attempt every 100 steps. During final STeUS simulations, the weights were stable and all states were visited with similar occupancies (Fig. S3). Only simulation frames at the ground temperature *T*_0_ were used to collect the US histograms and, hence, to compute the PMF.

### Replica-exchange umbrella sampling: REUS

The CV *z*, number of umbrella windows, initial configurations, restraints used during production, total simulation time, and US force constant were chosen as during standalone US. To choose the spacing between the 144 umbrella windows along *z* we ran a series of 60 ps simulations and optimized the spacing to reach an acceptance probability for exchanges between neighboring windows in the range of 0.25% to 0.47%. During our optimization process, we assumed a linear relation between the *z*-spacing and the rejection probability within the tested range of window spacings (Fig. S4). Then, we selected a *z*-spacing between windows with an expected exchange rejection probability of 0.64%. Because running 144 umbrella windows in parallel would require the simultaneous allocation of many compute nodes, we grouped our umbrella windows into eleven subsets along the *z*-range, and we allowed exchanges only within the subsets. To guarantee good overlap between umbrella windows at the ends of the subsets, neighboring subsets overlapped by two umbrella windows (Figs. S5 and S6). Because the constriction region of OprO at [−1, 1] nm was subject to increased sampling problems due to extensive contacts between the porin and fosmidomycin, we used more replicas for the subset close to this region. Hence, the subset centered at *z* = 0 nm was composed of 24 windows, while all other subsets were composed of 16 windows except for the two subsets including the *z*-range extrema which were composed of only eight windows. The average exchange probabilities of our production REUS were consistent with our optimization procedure, *i.e.*, nearly all exchange probabilities were within the [0.25%, 0.47%] range with only few exceptions (Figs. S5 and S6).

### Restraints used during initial pulling and during umbrella sampling

During the initial pulling simulation (steered MD) and during US, we applied on the fosmidomycin an orientational flat-bottomed restraint as well as a cylindrical flat-bottomed restraint. During the initial pulling and equilibration only, we applied in addition position restraints on protein atoms, as described and rationalized in the following sections.

#### Flat-bottomed orientation restraints

The angle between the vector connecting the two ends of the drug and the *z*-axis was restrained with a flat-bottomed potential within [-45°, 45°], using a force constant of 7878 kJ mol*^−^*^1^ rad*^−^*^2^ outside of this interval. The angle restrains were used to maintain fosmidomycin either in orientation 1 or in orientation 2 as defined in Fig. 2C. The orientation restraint was strictly required to obtain converged PMFs because sampling of a orientational flip is extremely rare in the narrow pore (if not impossible). As such, we computed two PMFs, one for each orientation, which may be combined a posteriori if needed.

#### Flat-bottomed cylindrical restraint along the pore axis

In addition, we applied a flat-bottom potential to restrain the antibiotic within a cylinder of radius 1 nm centered on the COM of C*_α_* atoms close to the cavity with a force constant of 1000 kJ mol*^−^*^1^ nm*^−^*^2^ acting outside of the cylinder in the direction of the cylinder axis (Fig. 2B). The cylindrical flatbottomed potential ensured that the solute would not “miss” the entrance to the pore, while the flat-bottomed region was large enough to ensure that the solute did not feel the potential if located in the constriction regions of the pore. Furthermore, the cylindrical flat-bottomed potentials serves to obtain a bulk state with well-defined entropy and free energy. Although not considered in this study, such well-defined bulk free energy would be needed to compute the overall permeability of a membrane with a given lateral concentration of porins.

#### Protein restraints during initial steered MD

To mitigate non-equilibrium effects during pulling simulations, we used position restraints on heavy atoms of the protein backbone and side chains with force constants of 1000 kJ mol*^−^*^1^ nm*^−^*^2^ and 100 kJ mol*^−^*^1^ nm*^−^*^2^ respectively. These restraints excluded that the protein would be distorted during pulling simulations, which would lead to hysteresis problems, i.e., to different PMFs obtained from pulling along PP-to-EC and for EC-to-PP directions. These restraints were gradually decreased during equilibration (see above) and fully removed during production simulations.

## Results

### Permeation of fosmidomycin in orientation 1 with standalone US

In the work from Golla *et al.*,^16^ PMFs of the permeation of fosmidomycin through OprO computed with standalone US exhibited hysteresis effects. To test how REUS, STeUS, and US-HREX improve the PMFs relative to standalone US, we first computed reference PMFs with standalone US. As CV, we used the *z*-component of the distance vector between the center of mass of OprO C*_α_* atoms close to the porin lumen (referred to as pCOM) and the center of mass of fosmidomycin phosphoryl group and amine group (referred to as fCOM) (Fig. 2B). Henceforth, we refer to this CV as *z*. We reduced the accessible configurational space by (i) restraining the orientation of the antibiotic relative to the *z*-axis within ±45°(Fig. 2C), and (ii) by applying a flat-bottom potential restraining the distance between the pCOM and the fCOM projected onto the *xy*-plane below 1 nm (Fig. 2B, see Methods). By using an orientational restraint, fosmidomycin could enter the porin from the EC either by presenting its amine group first (orientation 1) or its phosphoryl group first (orientation 2) (Fig. 2C). For standalone US we investigated only orientation 1.

Because the free energy is a state function, PMFs computed with umbrella sampling should not depend on the path taken by the initial pulling simulations^43, 44^ used to generate the initial configurations. A sensitive test for the convergence of PMFs computed with umbrella sampling is to compare PMFs obtained with initial configurations generated from “forward” and “reverse” pulling simulations, here corresponding to initial pulling in EC-to-PP and PP-to-EC directions, respectively. Figure 3A/B presents PMFs obtained from EC-to-PP or PP-to-EC pulling simulations, where the PMFs were defined to zero at the largest *z*-position. Each panel shows four PMFs, based on two independent pulling simulations (repl. 1 and repl. 2), while each 200 ns US window was furthermore split into two time-blocks of 0– 100 ns and 100–200 ns. The average PMFs for EC-to-PP and PP-to-EC directions are shown in Fig. 3C together with two standard errors as shaded areas, revealing 95% confidence intervals up to 7.5 kcal mol*^−^*^1^ or up to 5.0 kcal mol*^−^*^1^ for the the EC-to-PP or PP-to-EC directions, respectively, demonstrating major statistical uncertainties. In addition, the free energy difference between the PP and EC endpoints strongly differs between the EC-to-PP and PP-to-EC PMFs, demonstrating a major hysteresis problem. Notably, without the use of restraints during US and initial pulling (see Methods), these hysteresis problems would likely be even more pronounced.

**Figure 3:**
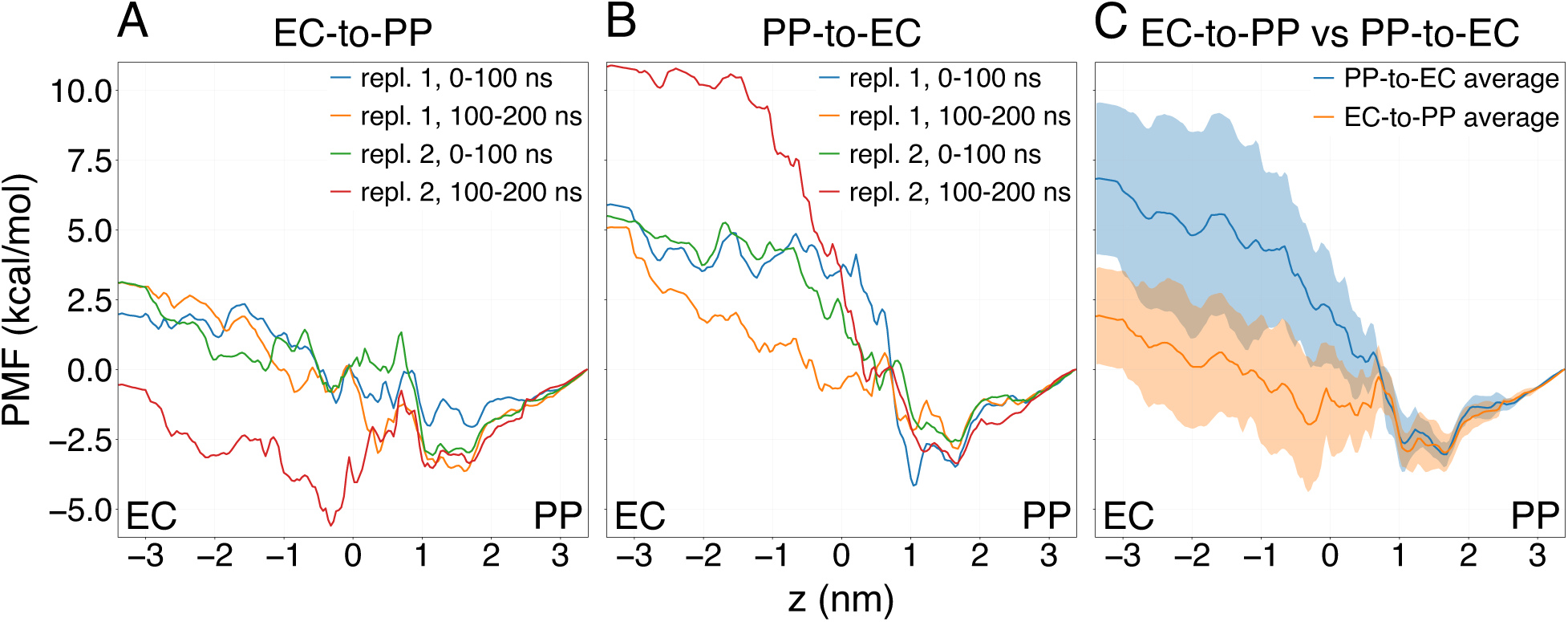
PMFs of fosmidomycin permeation in orientation 1 with standalone US obtained after (A) EC-to-PP pulling or (B) PP-to-EC pulling. Two independent pulling simulation replicates were carried out for each direction and used to obtain initial frames for US. For each replicate, umbrella windows were binned into two time-blocks of 0–100 ns and 100– 200 ns, yielding a total of four PMFs for each permeation direction (blue, orange, red, and red lines). (C) Average PMF for each direction. Confidence intervals (shaded areas) represent two standard errors. Evidently, PMFs from standalone US are poorly converged and subject to major hysteresis problems. {us}

To shed light on the molecular interactions underlying the hysteresis, we inspected the umbrella potential energy over time in different umbrella windows to detect high variation of the bias potential. We observed a bias potential peak in the umbrella window centered at *z* = −0.007 nm (Fig. 4A); this peak correlates with the presence of a water molecule trapped between fosmidomycin and the protein (Fig. 4B, red circle). Thus, solvent degrees of freedom likely contributed to the observed hysteresis effects. In line with the work from Golla *et al.*, our results confirm that using *z* as a collective variable with standalone umbrella sampling is not sufficient to sample all relevant degrees of freedom involved in fosmidomycin permeation through OprO using 200 ns per US window, even in the presence of several restraints.

**Figure 4:**
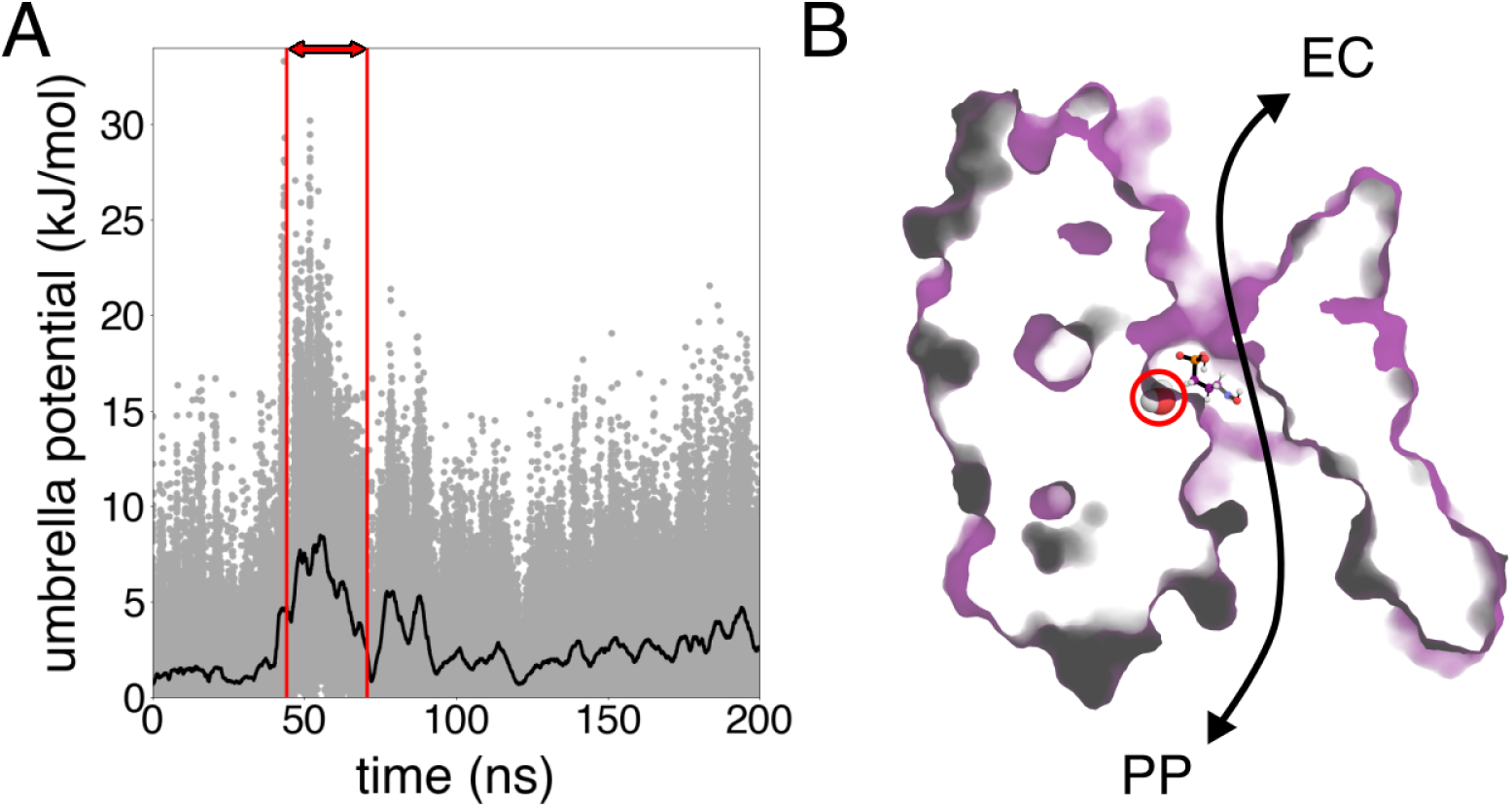
On the influence of water on poor convergence during standalone US. (A) Bias potential energy of umbrella window centered at *z* = −0.007 nm vs. simulation time (grey dots) and smoothed with the scipy module uniform_filter1d^45^ with a filter size of 5000 points (black line). (B) Representative snapshot during simulation time between 45 ns and 71 ns (red box in panel A). The water molecule trapped between fosmidomycin (represented as balls and sticks) and the porin (represented as a surface) is highlighted with a red circle. The double-headed black arrow sketches the OprO lumen. {bias_snap}

### Comparison of US flavors for fosmidomycin in orientation 1: US-HREX, STeUS, and REUS

To overcome the sampling and hysteresis problems with standalone US (previous paragraph), we tested three improved US flavors, namely US-HREX, STeUS, and REUS. To apply US-HREX in practice, it is critical to understand the molecular interactions underlying the free energy barriers along the orthogonal degrees of freedom. Here, because cationic residues inside the OprO lumen strongly interact with fosmidomycin,^16^ we chose to scale positive charges of the porin and negative charges within the phosphoryl group of fosmidomycin by the *λ*-parameter along 24 *λ*-replicas per US window (see Methods for details).

Increasing the temperature is another mean to improve sampling. This principle is exploited in the simulated tempering framework, where a Metropolis criterion is used to accept or reject steps along a pre-defined temperature ladder within a single simulation. In higher temperature states, the probability to cross enthalpic barriers together with exchanges with low temperature states will improve configurational sampling at all temperatures including the base temperature. Combining simulated tempering with umbrella sampling in STeUS has been highly successful during PMF calculations for drug permeation across a lipid membrane.^23^ In this study, simulated tempering was applied in each umbrella window with temperatures ranging from 300 K to 348 K with 4 K-steps, and only data acquired at the base temperature was used to compute the PMFs (see Methods for details).

REUS (also referred as bias-exchange US) exploits the fact that neighboring umbrella windows can explore different regions of phase space and, therefore, by exchanging configurations between windows according to a Metropolis criterion,^46^ improved sampling of relevant degrees of freedom orthogonal to the CV is expected. In REUS, it is common practice to permit configuration exchanges along the whole CV-space. In this study, in contrast, we permitted exchanges only between windows within subsets of *z* to reduce the amount of computational resources needed simultaneously (see Methods).

PMFs of the fosmidomycin permeation through OprO obtained with the three aforementioned methods are shown in Fig. 5. The PMFs obtained with US-HREX exhibit a steep rise at *z* ≈1 nm, suggesting that these PMFs suffer from hysteresis problems (Fig. 5A/B). In addition, the EC-to-PP and PP-to-EC PMFs from US-HREX strongly differ within the region |*z*| *<* 1 nm, and the averaged PMFs exhibit large statistical errors (Fig. 5C). Hence, US-HREX hardly improved the sampling relative to standalone US. Compared to the PMFs from US-HREX, the PMFs from STeUS are slightly more converged, as evident from the slightly reduced uncertainties (Fig. 5F) and from the absence of a spurious rise at *z* ≈1 nm (Fig. 5D/E). Nevertheless, the major statistical uncertainties remain with STeUS as well as a major hysteresis between EC-to-PP and PP-to-EC directions.

**Figure 5:**
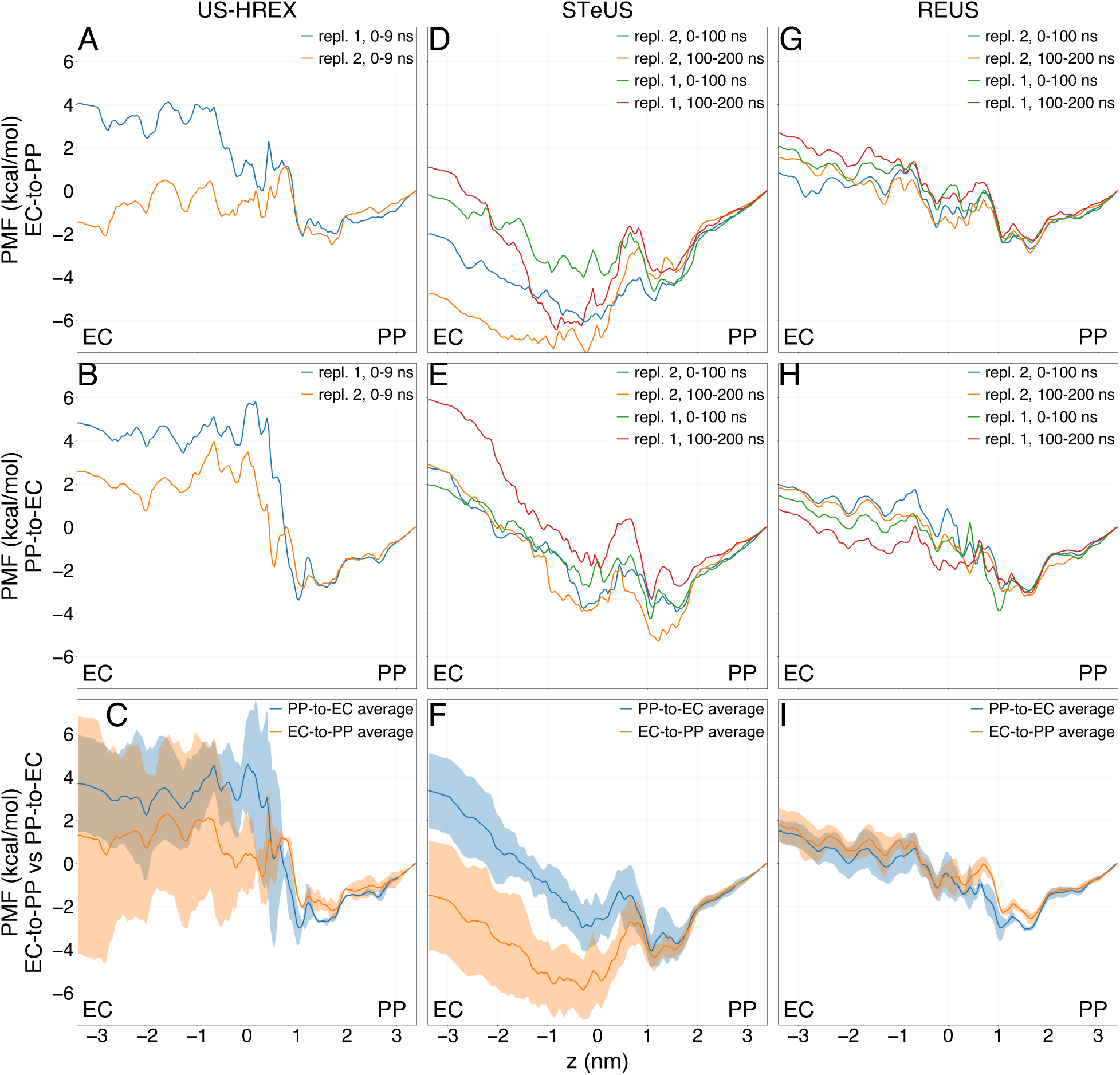
PMFs of fosmidomycin permeation in orientation 1 obtained with US-HREX (A– C), STeUS (D–F) or REUS (G–I). EC-to-PP (A/D/G) and PP-to-EC PMFs (B/E/H) were computed with initial conformations obtained from EC-to-PP and PP-to-EC pulling simulations, respectively. For each EC-to-PP and PP-to-EC setup, two independent pulling simulations were carried out: replicate 1 and 2. For STeUS and REUS, umbrella windows were binned into time-blocks 0–100 ns and 100–200 ns. (C/F/I) Averages of EC-to-PP (orange) and PP-to-EC (blue) PMFs by combining all respective umbrella windows. Confidence intervals (shaded areas) estimated from independent PMFs represent two standard errors. {reus_steus_

To understand why PMFs of the permeation of fosmidomycin through OprO computed with STeUS display hysteresis problems, we visually inspected the trajectories. Accordingly, we noticed that fosmidomycin has been trapped in a pocket at the EC entrance of OprO in the umbrella window centered at *z* = −1.63 nm (Fig. 6). The transition towards this pocket occurred after ∼100 ns, while fosmidomycin remained in this pocket for the remaining 100 ns, irrespective of a uniform coverage of all temperature states. Hence, simulated tempering did not help fosmidomycin to escape from this pocket within simulation time. To further test the stability of this unexpected state, we ran three free simulations of 100 ns each starting from configurations extracted at *t* =130 ns, *t* =160 ns and *t* =200 ns of this US window. Fosmidomycin did not escape the pocket in any of these simulations. Besides the population of this unexpected state that does not contribute to successful permeation, we furthermore observed strongly increased flexibility of protein loops relative to the other US flavors as consequence of populations of higher-temperature states during simulated tempering. These results suggest that, for simulating permeation across OprO, the use of simulated tempering increases the risks of (i) populating non-productive conformation and (ii) of partly unfolding the protein.

**Figure 6:**
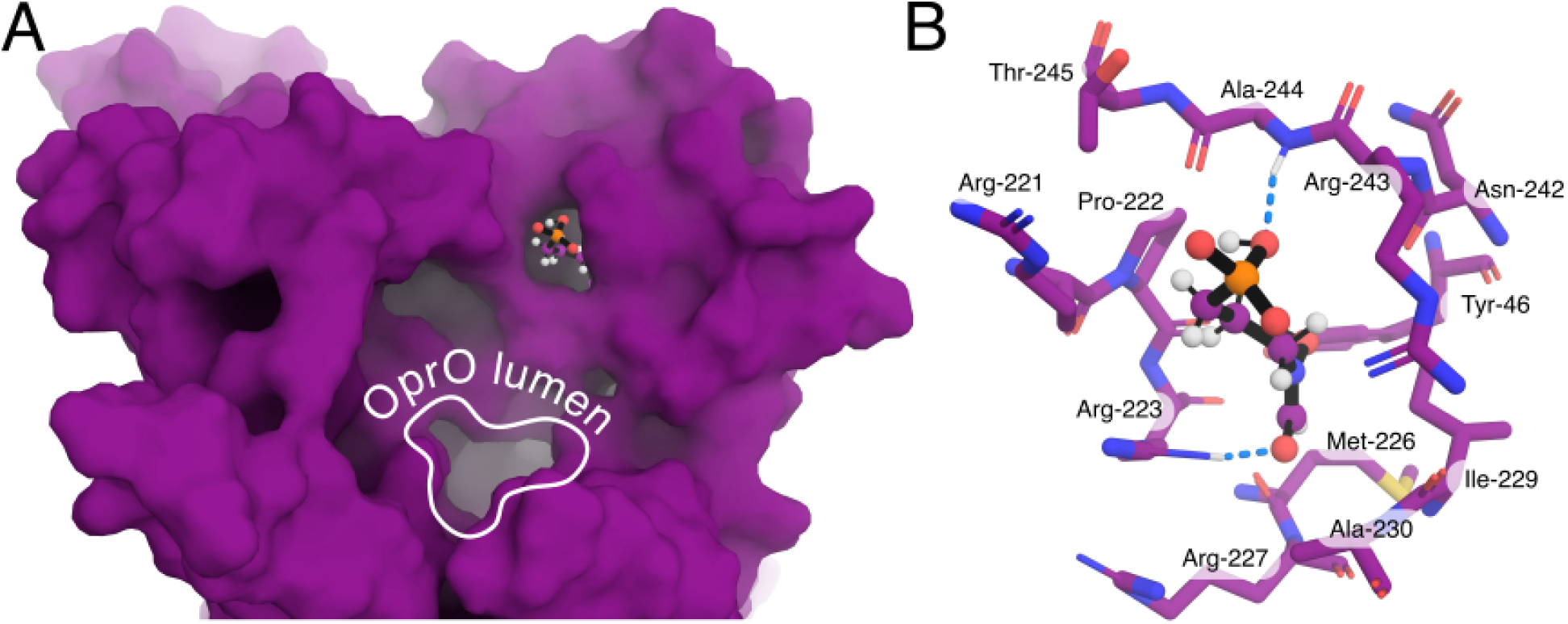
(A) Snapshot of the umbrella window centered at *z* = − 1.63 nm from STeUS, revealing fosmidomycin trapped in a pocket near the EC entrance. OprO is represented as a purple surface, and fosmidomycin as balls and sticks. (B) Focus on fosmidomycin and protein residues of the pocket. Blue dotted lines indicate fosmidomycin–protein hydrogen bonds. {st_stuck}

In sharp contrast to the PMFs obtained with US-HREX or STeUS, PMFs obtained with REUS were highly converged and exhibited virtually no hysteresis problems (Fig. 5G–I). Along the complete CV, the EC-to-PP and PP-to-EC PMFs revealed excellent agreement with a maximum deviation of ∼1 kcal/mol (Fig. 5I), while the 95% confidence intervals were smaller or close to 1 kcal/mol (Fig. 5I, shaded areas), by far lower as compared to the confidence intervals obtained with US-HREX, STeUS, or standalone US (Fig. 3C and 5C/F and ). Taken together, among the four US flavors considered in this study, REUS provides by far the best converged PMFs of fosmidomycin permeation across OprO.

The agreement of EC-to-PP PMFs with PP-to-EC PMFs obtained from REUS justifies the averaging of these PMFs to further reduced the statistical uncertainties. By averaging all EC-to-PP and PP-to-EC PMFs computed with REUS, we obtained PMFs for fosmidomycin permeation in orientation 1 with 95% confidence intervals of approximately 0.5 kcal/mol (Fig. 7, blue curve). The free energy minimum of the PMF in orientation 1 at *z* = 1.6 nm corresponds to a wider region of the pore, where fosmidomycin may adopt different sets of interactions with Arg^34^, Lys^321^, or Lys^388^ (Fig. 7, right panel).

**Figure 7:**
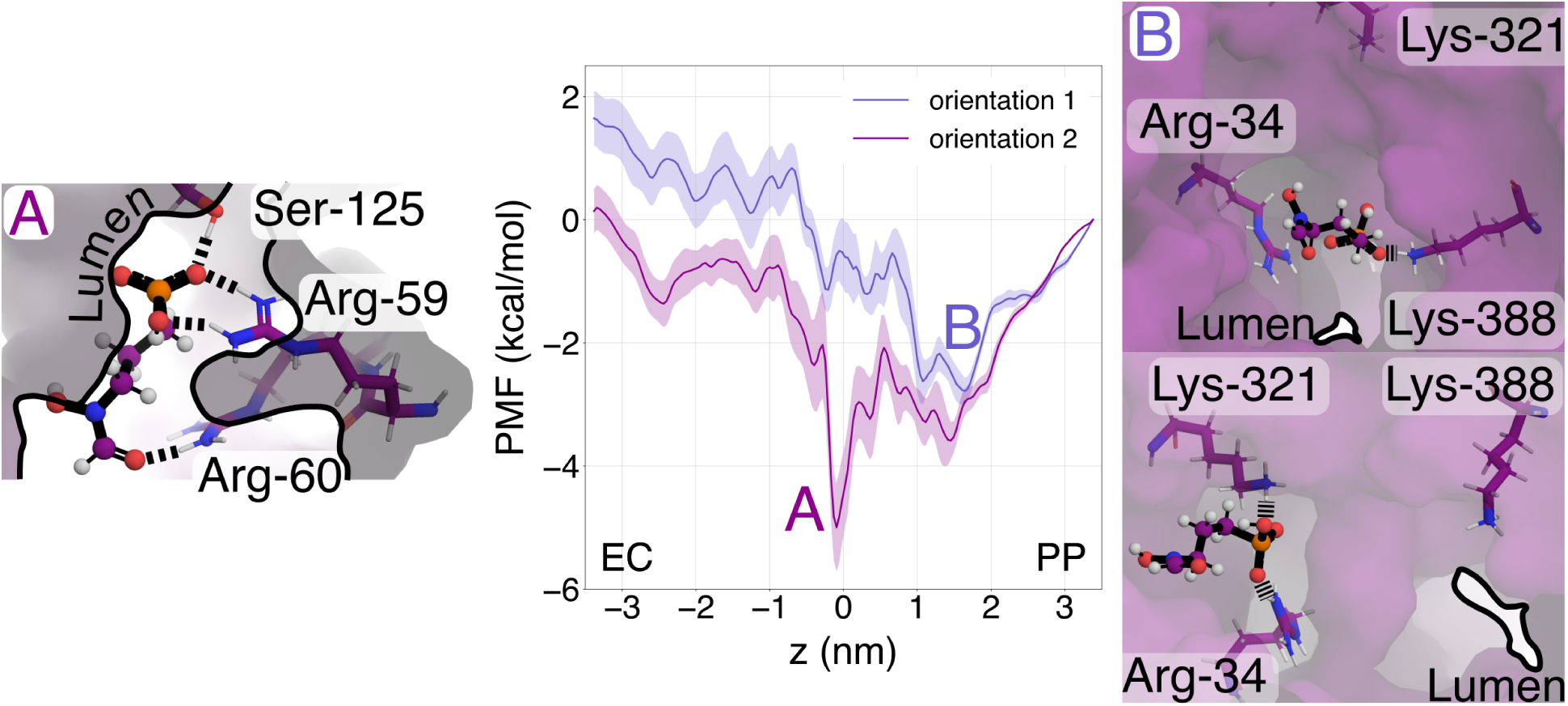
Averaged PMFs of fosmidomycin permeation in orientation 1 (dark blue) or orientation 2 (magenta) computed with REUS, after combining EC-to-PP and PP-to-EC calculations. Example simulation snapshots of conformation of the free energy minima for orientation 1 and 2 are depicted in panels B and A, respectively. Confidence intervals (shaded areas) represent two standard errors. {or1_or2}

### Permeation of fosmidomycin in orientation 2 with REUS

The PMFs discussed in the sections above correspond to orientation 1 of fosmidomycin with the phosphate moiety pointing towards the extracellular side (Fig. 2C). We proceeded to test whether REUS likewise provides converged PMFs for fosmidomycin in orientation 2, initially using the same computational effort as used for orientation 1. However, unlike the PMFs for orientation 1 that were largely overlapping among the EC-to-PP and PP-to-EC directions (Fig. 5G–I), the PMFs for orientation 2 overlapped only in the region between 0.5 nm and 3.5 nm, revealing increased hysteresis problems (Fig. 8A). In our implementation of REUS, exchanges between neighboring US windows were only allowed within subsets of the *z*-space. Therefore, we hypothesized that sampling could be improved in regions of *z*-space that exhibit the largest discrepancies between EC-to-PP and PP-to-EC PMFs by (i) running additional simulations and (ii) by increasing the number of windows that may exchange configurations.

**Figure 8:**
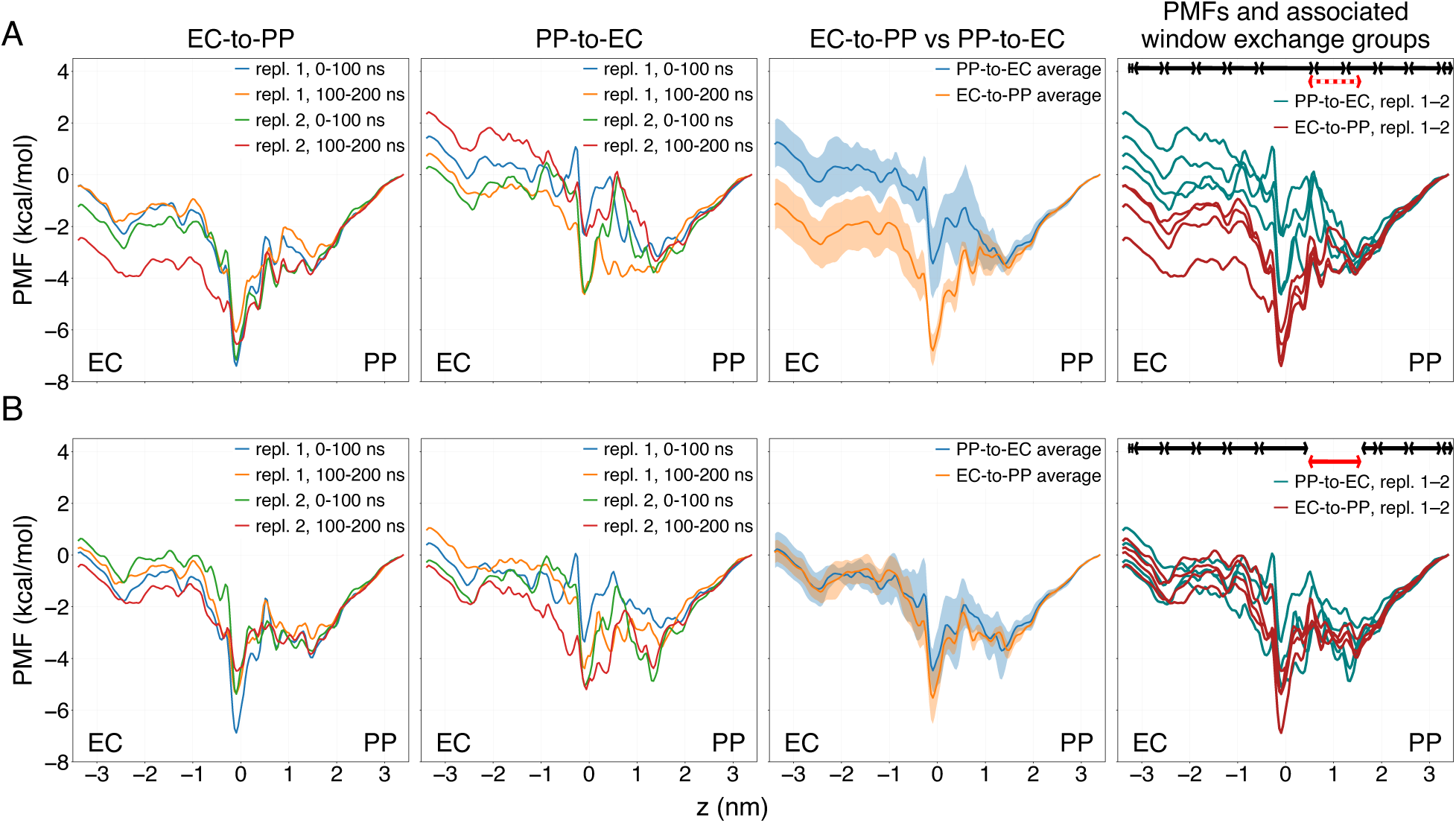
PMFs of fosmidomycin permeation in orientation 2 with REUS. (A) PMFs of the permeation process with sub-optimal choice of window subsets within which exchanges are allowed. EC-to-PP and PP-to-EC PMFs refer to REUS setups started with initial conformations obtained from EC-to-PP and PP-to-EC pulling simulations, respectively (two left columns, see panel headings). For each setup, two independent pulling simulations were carried out: replicate 1 and 2. Umbrella windows were binned into time-blocks 0–100 ns and 100–200 ns, yielding two PMFs for each direction and orientation (see legends). Third column: average EC-to-PP (orange) PP-to-EC (blue) PMFs, with confidence intervals representing two standard errors. Right column: US windows within *z*-subsets delimited by the dark arrows were carried out in parallel and, therefore, were allowed to exchange configurations. Region within the red-dotted arrow shows a high discrepancy between EC-to-PP and PP-to-EC PMFs, indicative of limited sampling. (B) PMFs of the permeation process with an optimized choice of window subsets within which exchanges are allowed. Descriptions of each column are identical to (A), except that data from a new REUS batch in the region *z*=[0.5, 3.5] nm (solid red arrow region in the fourth column) were used to obtain the PMFs. {reus_or2}

By overlaying all PMFs, we observed that the shape of the EC-to-PP and PP-to-EC PMFs largely agreed except in the region *z* ∈ [0.45 nm, 1.54 nm] (Fig. 8A, last column). Thus, we carried out an additional batch of REUS within this *z*-region for each replicate (red bar in Fig. 8B, fourth column), and we allowed exchanges between all windows within this batch. By computing new PMFs from the new REUS batch together the previous simulations, we obtained PMFs with favorable agreement between EC-to-PP and PP-to-EC directions, hence revealing the absence of hysteresis problems (Fig. 8B). The 95% confidence intervals were ≤1.5 kcal/mol throughout the CV for both PP-to-EC and EC-to-PP PMFs (Fig. 8B, 3^rd^ column). Hence, we obtained well-converged PMFs also in orientation 2 by (i) identifying the undersampled region via comparing PMFs from forward and reverse pulling, and (ii) augmenting the sampling in the undersampled regions.

By averaging all EC-to-PP and PP-to-EC PMF replicates in Fig. 8B, we obtained the PMF with 95% confidence intervals below 1 kcal/mol for orientation 2 (Fig. 7, magenta curve). The PMF for orientation 2 reveals a marked free energy minimum at the channel center at *z* = −0.1 nm, where fosmidomycin forms simultaneous hydrogen bonds with Arg^59^, Arg^60^, and Ser^125^ (Fig. 7, left panel).

## Discussion

We compared four different US flavors for studying the permeation of fosmidomycin through the OprO porin: standalone US, REUS, STeUS, and US-HREX. Among these methods, REUS revealed by far the best converged PMFs. For a given fosmidomycin orientation, using a total simulation time of approximately 65µs, REUS achieved PMFs that were converged with 95% confidence intervals (two standard errors) well below one kilocalorie per mole. Although the computational effort for obtaining the converged PMF is considerable, we expect that, with ever increasing computer power, simulations as presented here may soon become routine for studying antibiotic uptake by outer membrane porins.

Our key assessment of convergence was given by the comparison of PMFs obtained after constant-velocity pulling simulations in forward or reverse directions, here corresponding to PP-to-EC or EC-to-PP directions, respectively. Because PMF calculations frequently suffer from hysteresis problems, comparing PMFs obtained after forward and reverse pulling provides a highly sensitive test for convergence.^47–49^ In addition, for each pulling direction (PP-to-EC or EC-to-PP), we computed PMFs from two independent set of US windows initiated from two independent pulling simulations. Comparing the latter PMFs provides an additional quality test, but it may not reveal hysteresis problems that are, in our experience, a major matter of concern during US simulations of biomolecules. An alternative strategy of estimating statistical errors may be given by splitting the US windows into time blocks, followed by comparison of the PMFs obtained from each time block. However, because autocorrelation times are typically unknown and may exceed the simulation time of US windows, the PMFs obtained from time blocks may not be independent and statistical errors may be severely underestimated. Hence, we suggest that the comparison of PMFs obtained after pulling in forward and reverse direction should become routine when reporting PMFs from US simulations of biomolecular simulations.

Poor conformational sampling is a consequence of long autocorrelation times owing to long-living conformational arrangements of solute, protein, and channel-bound water molecules. Since both HREX and simulated tempering have been used successfully to enhance the conformational sampling of protein or membrane systems,^23, 50–52^ we anticipated that augmenting US with HREX (US-HREX) or with simulated tempering (STeUS) would accelerate the converge of PMF calculations of fosmidomycin permeation. However, US-HREX and STeUS provided only a small sampling benefit (if any) as compared do standalone US. The benefit of US-HREX is furthermore reduced owing to the requirement of simulating parallel replicas of each US window. Moreover, the use of US-HREX is complicated by the fact it requires the selection of a set of interactions to be scaled along the *λ*-variable, which is *a priori* far from obvious. In our implementation of US-HREX, we scaled the charges of fosmidomycin and cationic residues with the aim to reduce the life time of salt bridges at higher *λ*-states and, thereby, to reduce autocorrelation times. Such choice may provide room for further optimization, for instance by restricting the scaling to cationic residues along the pore lumen. However, since US-HREX was hardly successful and computationally expensive, we did not further optimize the selection of scaled interactions.

We recently observed an at least five-fold enhanced sampling by STeUS relative to standalone US for drug permeation across a lipid membrane.^23^ For fosmidomycin permeation across OprO, in contrast, STeUS provided only a small benefit, suggesting that higher temperatures reduced autocorrelation times only moderately. A disadvantage of the simulating tempering simulations as observe here may be an increased risks of visiting conformations that do not contribute to the permeation process (Fig. 4), or of perturbing the protein structure at higher temperatures. Such risk may not apply in disordered systems such as lipid membranes. Hence, additional simulations will be required to clarify which systems benefit from simulated tempering during US.

According to our REUS protocol, we split our CV-space into several subsets of eight to 24 neighboring US windows, allowing exchanges of configurations only within a subset. Compared to more common REUS setups that allow exchanges among all windows within a single simulation batch, our protocol greatly reduces the number simultaneously allocated compute nodes, hence allowing efficient use of compute clusters without longer waiting times and the use of REUS simulations on commodity clusters. Furthermore, such protocol readily allows dedicating more resources and longer simulation times to CV regions that are critical for sampling.

A critical requirement for obtaining converged PMFs with REUS was the use of several types of restraints. (i) We restrained the orientation of fosmidomycin relative to the *z*-axis within ±45°. The orientational restraint avoids long autocorrelation times owing to slow conformational sampling of the fosmidomycin orientation in the pore lumen. Indeed, during an early stage of this project, we obtained poorly converging PMFs in REUS simulations without orientation restraints. (ii) A flat-bottomed cylindrical restraint acting in lateral direction kept fosmidomycin near the pore axis. Such restraint excluded that the solute diffuses laterally in the complete *xy*-plane of the simulation box, which leads to sampling problems in US windows near the pore entrance. (iii) During initial constant-velocity pulling simulations of fosmidomycin across the pore, it was critical to restrain the positions of protein atoms. Without such restraints, the protein structure may be perturbed because the steered fosmidomycin may drag protein residues or distort geometric constriction sites. Because the equilibration time of US windows is typically insufficient to mitigate such structural perturbations, the perturbations would lead to undesired hysteresis problems.

Quantitative predictions of membrane permeability for antibiotics involve several additional challenges, which were not addressed in this study. If the porin carries out larger conformational transitions during antibiotic permeation, for instance involving flexible loops, additional sampling challenges emerge.^25^ Obtaining the permeability requires, apart from the PMF, also the calculation of the position-dependent diffusion coefficient.^53^ Because the anionic fosmidomycin binds to the cationic OprO lumen as shown by the free energy minima along the PMF (Fig. 7), fosmidomycin permeation may involve competitive binding with other anionic species such as phosphate ions, which were not simulated here. Furthermore, apart from overcoming sampling challenges addressed here, accurate permeability predictions rely on accurate force fields; however, considering that salt bridges are subject to uncertainty in biomolecular simulations, ^54^ careful force field validations will be critical for obtaining accurate PMFs for the permeation of ionic solutes such as fosmidomycin.

## Conclusions

We compared four different US flavors used to compute the PMF of permeation of the antibiotic fosmidomycin across the outer membrane porin OprO: standalone US, US augmented with HREX (US-HREX) or with simulated tempering (STeUS), as well as REUS. In contrast to PMFs obtained with standalone US, US-HREX, or STeUS, the PMFs obtained with REUS were well converged as shown by the absence of hysteresis between PMF calculations carried out in forward and reverse directions and by 95% confidence intervals below one kilocalorie per mole. The convergence of PMFs obtained with REUS relied on the use of several geometric restraints that helped the simulations to circumvent long autocorrelation times and to avoid structural perturbations during initial pulling simulations. We anticipate that the systematic comparison of US flavors as well as the protocol for obtaining converged PMFs of OprO permeation will be useful for studying antibiotic uptake over various porins in future studies.

## Supporting information

Supporting material for: Converging PMF calculations of antibiotic permeation across an outer membrane porin with sub-kilocalorie per mole accuracy

## Acknowledgement

We thank Ulrich Kleinekathöfer and Vinaya Kumar Golla for fruitful discussions, for pointing us to the challenges of converging PMF calculations of porin permeation, and for kindly sharing the simulation systems and forcefield parameters of OprO and fosmidomycin. This study was supported by the Deutsche Forschungsgemeinschaft (DFG, German Research Foundation) via grants SFB 860/A16 and INST 256/539-1

