## Supporting material for: Converging PMF calculations of antibiotic permeation across an outer membrane porin with sub-kilocalorie per mole accuracy for "Converging PMF calculations of antibiotic permeation across an outer membrane porin with sub-kilocalorie per mole accuracy"

A

| umbrella window centers (nm) | umbrella window centers (nm) | umbrella window centers (nm) | umbrella window centers (nm) | umbrella window centers (nm) | umbrella window centers (nm) | umbrella window centers (nm) | umbrella window centers (nm) | umbrella window centers (nm) | umbrella window centers (nm) |
|---|---|---|---|---|---|---|---|---|---|
| -3.35 | -2.64 | -1.95 | -1.23 | -0.54 | 0.21 | 0.89 | 1.59 | 2.28 | 2.98 |
| -3.31 | -2.60 | -1.91 | -1.18 | -0.48 | 0.26 | 0.94 | 1.63 | 2.33 | 3.02 |
| -3.26 | -2.55 | -1.86 | -1.12 | -0.41 | 0.30 | 0.99 | 1.68 | 2.38 | 3.07 |
| -3.21 | -2.51 | -1.81 | -1.08 | -0.34 | 0.35 | 1.04 | 1.72 | 2.42 | 3.11 |
| -3.16 | -2.46 | -1.75 | -1.06 | -0.32 | 0.38 | 1.09 | 1.77 | 2.47 | 3.16 |
| -3.12 | -2.41 | -1.71 | -1.01 | -0.28 | 0.43 | 1.13 | 1.82 | 2.51 | 3.20 |
| -3.07 | -2.37 | -1.66 | -0.95 | -0.22 | 0.48 | 1.17 | 1.86 | 2.56 | 3.25 |
| -3.02 | -2.32 | -1.62 | -0.90 | -0.17 | 0.52 | 1.21 | 1.91 | 2.61 | 3.29 |
| -2.97 | -2.27 | -1.57 | -0.86 | -0.14 | 0.57 | 1.26 | 1.96 | 2.65 | 3.34 |
| -2.93 | -2.23 | -1.52 | -0.81 | -0.09 | 0.61 | 1.31 | 2.01 | 2.70 | |
| -2.88 | -2.19 | -1.47 | -0.76 | -0.04 | 0.66 | 1.36 | 2.05 | 2.75 | |
| -2.83 | -2.14 | -1.41 | -0.71 | 0.00 | 0.70 | 1.41 | 2.10 | 2.79 | |
| -2.78 | -2.09 | -1.36 | -0.65 | 0.05 | 0.75 | 1.45 | 2.15 | 2.84 | |
| -2.74 | -2.04 | -1.32 | -0.61 | 0.10 | 0.80 | 1.49 | 2.19 | 2.88 | |
| -2.69 | -1.99 | -1.27 | -0.57 | 0.15 | 0.84 | 1.54 | 2.23 | 2.93 | |

B

| umbrella window centers (nm) | umbrella window centers (nm) | umbrella window centers (nm) | umbrella window centers (nm) | umbrella window centers (nm) | umbrella window centers (nm) | umbrella window centers (nm) | umbrella window centers (nm) | umbrella window centers (nm) | umbrella window centers (nm) |
|---|---|---|---|---|---|---|---|---|---|
| -3.40 | -2.67 | -1.96 | -1.26 | -0.56 | 0.12 | 0.85 | 1.58 | 2.27 | 2.97 |
| -3.35 | -2.62 | -1.91 | -1.22 | -0.51 | 0.19 | 0.89 | 1.62 | 2.32 | 3.01 |
| -3.30 | -2.58 | -1.87 | -1.18 | -0.47 | 0.25 | 0.94 | 1.67 | 2.37 | 3.06 |
| -3.25 | -2.53 | -1.82 | -1.13 | -0.42 | 0.29 | 1.00 | 1.72 | 2.41 | 3.10 |
| -3.20 | -2.48 | -1.77 | -1.09 | -0.37 | 0.33 | 1.05 | 1.76 | 2.46 | 3.15 |
| -3.16 | -2.43 | -1.73 | -1.04 | -0.34 | 0.38 | 1.10 | 1.81 | 2.51 | 3.19 |
| -3.11 | -2.38 | -1.69 | -0.98 | -0.30 | 0.43 | 1.14 | 1.86 | 2.55 | 3.24 |
| -3.06 | -2.34 | -1.64 | -0.94 | -0.25 | 0.47 | 1.19 | 1.91 | 2.60 | 3.29 |
| -3.01 | -2.30 | -1.60 | -0.89 | -0.20 | 0.52 | 1.24 | 1.95 | 2.64 | 3.34 |
| -2.96 | -2.24 | -1.56 | -0.85 | -0.15 | 0.56 | 1.28 | 2.00 | 2.69 | |
| -2.91 | -2.20 | -1.50 | -0.80 | -0.10 | 0.61 | 1.33 | 2.04 | 2.74 | |
| -2.87 | -2.15 | -1.46 | -0.74 | -0.06 | 0.66 | 1.39 | 2.08 | 2.79 | |
| -2.82 | -2.10 | -1.41 | -0.70 | -0.01 | 0.71 | 1.44 | 2.13 | 2.83 | |
| -2.77 | -2.05 | -1.36 | -0.66 | 0.03 | 0.77 | 1.48 | 2.17 | 2.87 | |
| -2.72 | -2.00 | -1.31 | -0.62 | 0.08 | 0.81 | 1.53 | 2.22 | 2.92 | |

C

| umbrella window centers (nm) | umbrella window centers (nm) | umbrella window centers (nm) | umbrella window centers (nm) | umbrella window centers (nm) | umbrella window centers (nm) | umbrella window centers (nm) | umbrella window centers (nm) | umbrella window centers (nm) | umbrella window centers (nm) |
|---|---|---|---|---|---|---|---|---|---|
| -3.45 | -2.72 | -2.01 | -1.31 | -0.59 | 0.10 | 0.81 | 1.52 | 2.22 | 2.93 |
| -3.40 | -2.67 | -1.97 | -1.26 | -0.55 | 0.14 | 0.86 | 1.57 | 2.27 | 2.98 |
| -3.35 | -2.62 | -1.92 | -1.21 | -0.50 | 0.20 | 0.90 | 1.62 | 2.32 | 3.02 |
| -3.30 | -2.58 | -1.87 | -1.17 | -0.45 | 0.27 | 0.95 | 1.66 | 2.37 | 3.07 |
| -3.25 | -2.53 | -1.82 | -1.12 | -0.40 | 0.33 | 0.99 | 1.71 | 2.42 | 3.12 |
| -3.20 | -2.48 | -1.77 | -1.07 | -0.36 | 0.37 | 1.04 | 1.75 | 2.46 | 3.16 |
| -3.16 | -2.43 | -1.72 | -1.03 | -0.32 | 0.40 | 1.09 | 1.80 | 2.50 | 3.21 |
| -3.11 | -2.38 | -1.68 | -0.97 | -0.27 | 0.44 | 1.15 | 1.84 | 2.55 | 3.26 |
| -3.06 | -2.34 | -1.63 | -0.92 | -0.22 | 0.49 | 1.20 | 1.89 | 2.60 | 3.31 |
| -3.01 | -2.29 | -1.59 | -0.88 | -0.17 | 0.54 | 1.24 | 1.94 | 2.65 | |
| -2.96 | -2.24 | -1.54 | -0.83 | -0.13 | 0.57 | 1.29 | 1.99 | 2.70 | |
| -2.91 | -2.20 | -1.49 | -0.79 | -0.08 | 0.62 | 1.33 | 2.03 | 2.75 | |
| -2.87 | -2.15 | -1.44 | -0.74 | -0.03 | 0.67 | 1.38 | 2.08 | 2.79 | |
| -2.82 | -2.11 | -1.39 | -0.70 | 0.01 | 0.71 | 1.43 | 2.13 | 2.84 | |
| -2.77 | -2.06 | -1.36 | -0.64 | 0.05 | 0.76 | 1.48 | 2.17 | 2.88 | |

Figure S1: Umbrella window centers for EC-to-PP (A) and PP-to-EC (B) in orientation 1, and for both EC-to-PP and PP-to-EC in orientation 2 (C). The same window centers were used for all umbrella sampling flavor.

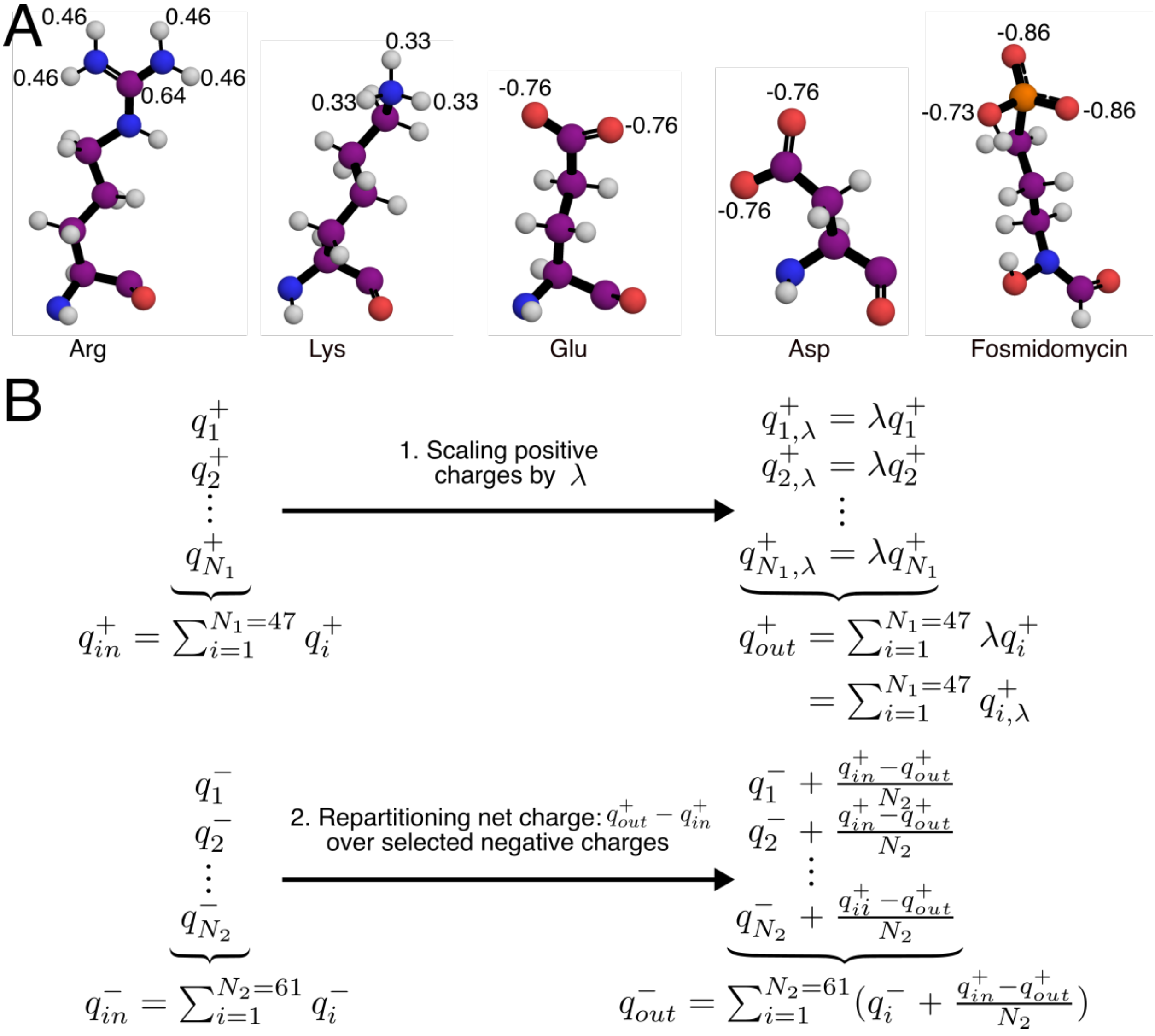


Figure S2: Hamiltonian replica-exchange protocol. (A) Residue types involved in charge scaling are depicted as balls and sticks, partial charges involved in charge scaling are written next to the respective atoms. (B) Scheme of our scaling protocol. First, positive charges $q^+_1$ through $q^+_{N_1}$ were scaled by a factor $\lambda$, leading to a net charge of $q^+_{out} - q^+_{in}$. To maintain charge neutrality, a set of $N_2$ negative charges was corrected by $-(q^+_{out} - q^+_{in})/N_2$.

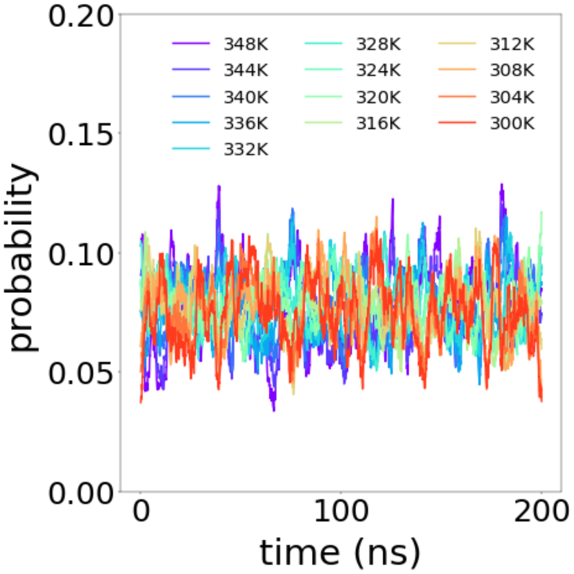


Figure S3: Temperature state probabilities versus time for window $z = 0\,\text{nm}$, revealing approximately uniform probabilities for all temperature states. Curves were smoothed with the Scipy module uniform_filter1d[1] with a filter size of 500 points.

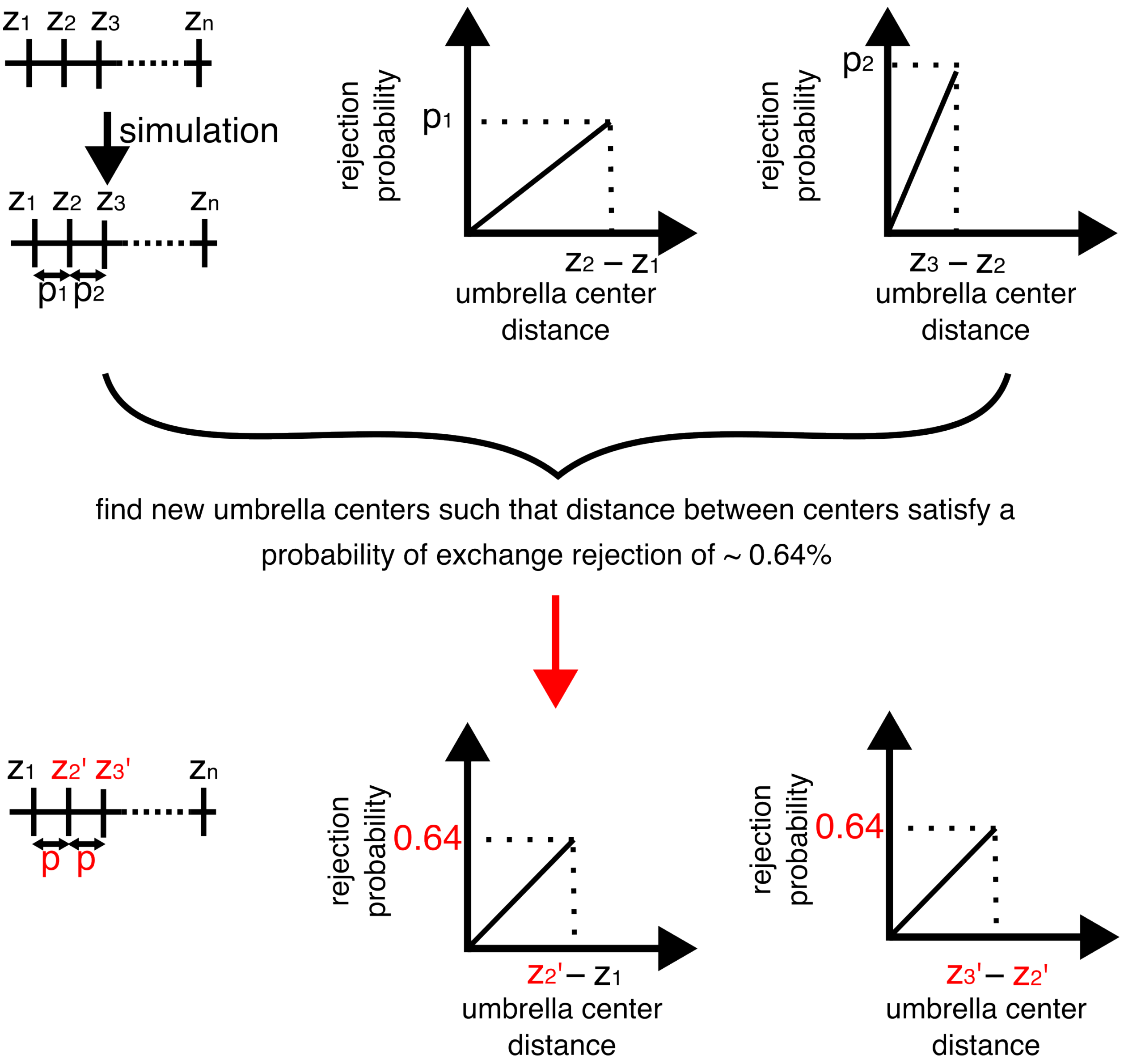


Figure S4: Protocol for optimizing the distance between neighboring umbrella windows for REUS. Series of simulations were carried out to compute average probabilities of exchange rejection between windows. From these probabilities, linear relationships between probabilities and distances between US windows were assumed. From this linear relationships, we selected centers of umbrella windows such that a rejection probability of approx. 0.64% was expected (acceptance probability of approx. 0.36%).

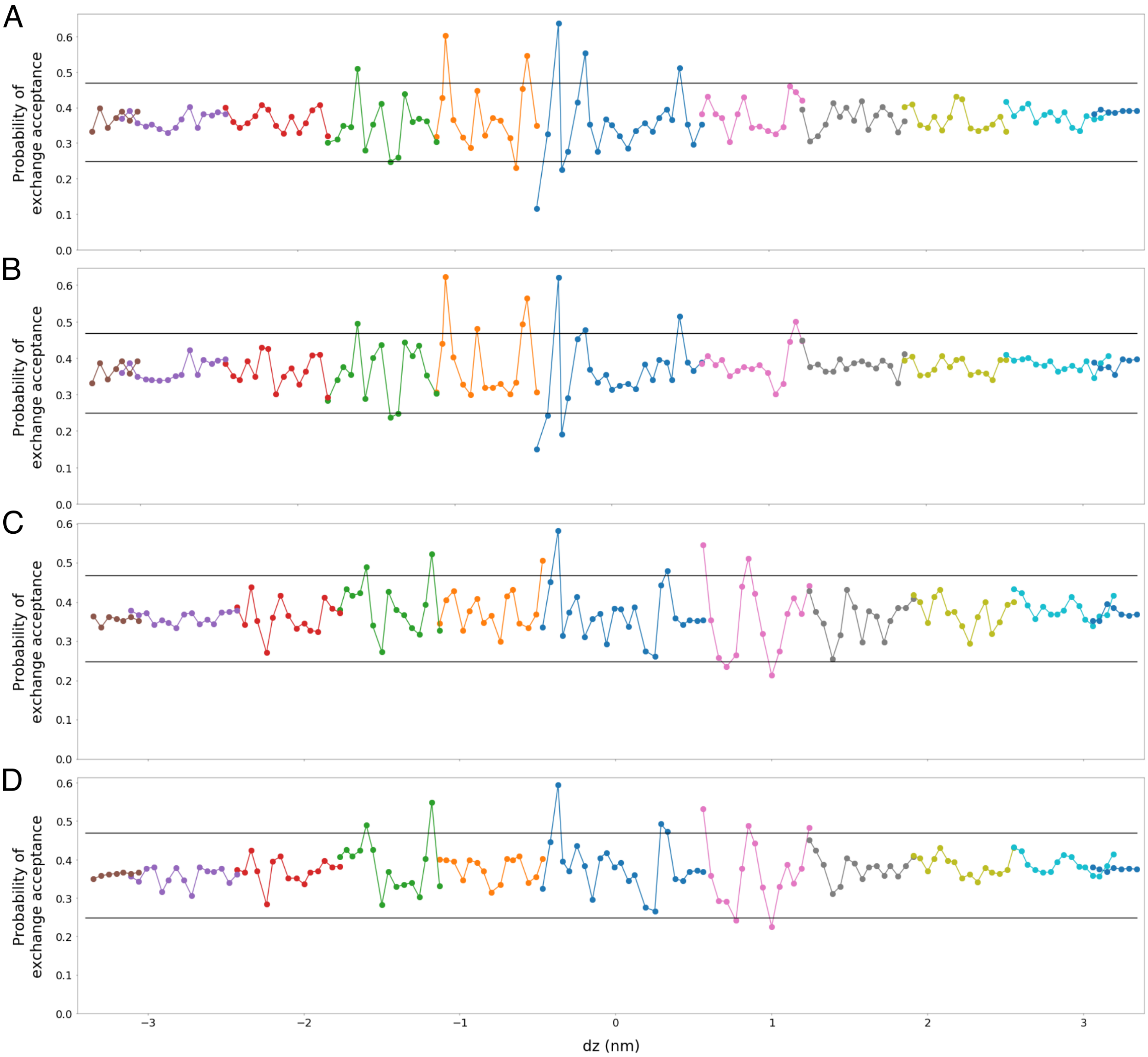


Figure S5: Average exchange probabilities between umbrella windows after 200 ns of production simulation for two forward replicates (A–B), and two reverse replicates (C–D) in orientation 1. Umbrella windows that were allowed to exchange configurations are shown with the same color. Each point shows the average exchange probability between the umbrella window centered at the respective $z$ value with the preceding umbrella window along the $z$-axis. Probability region between the two horizontal black lines indicates the target region used for optimizing the umbrella window spacing.

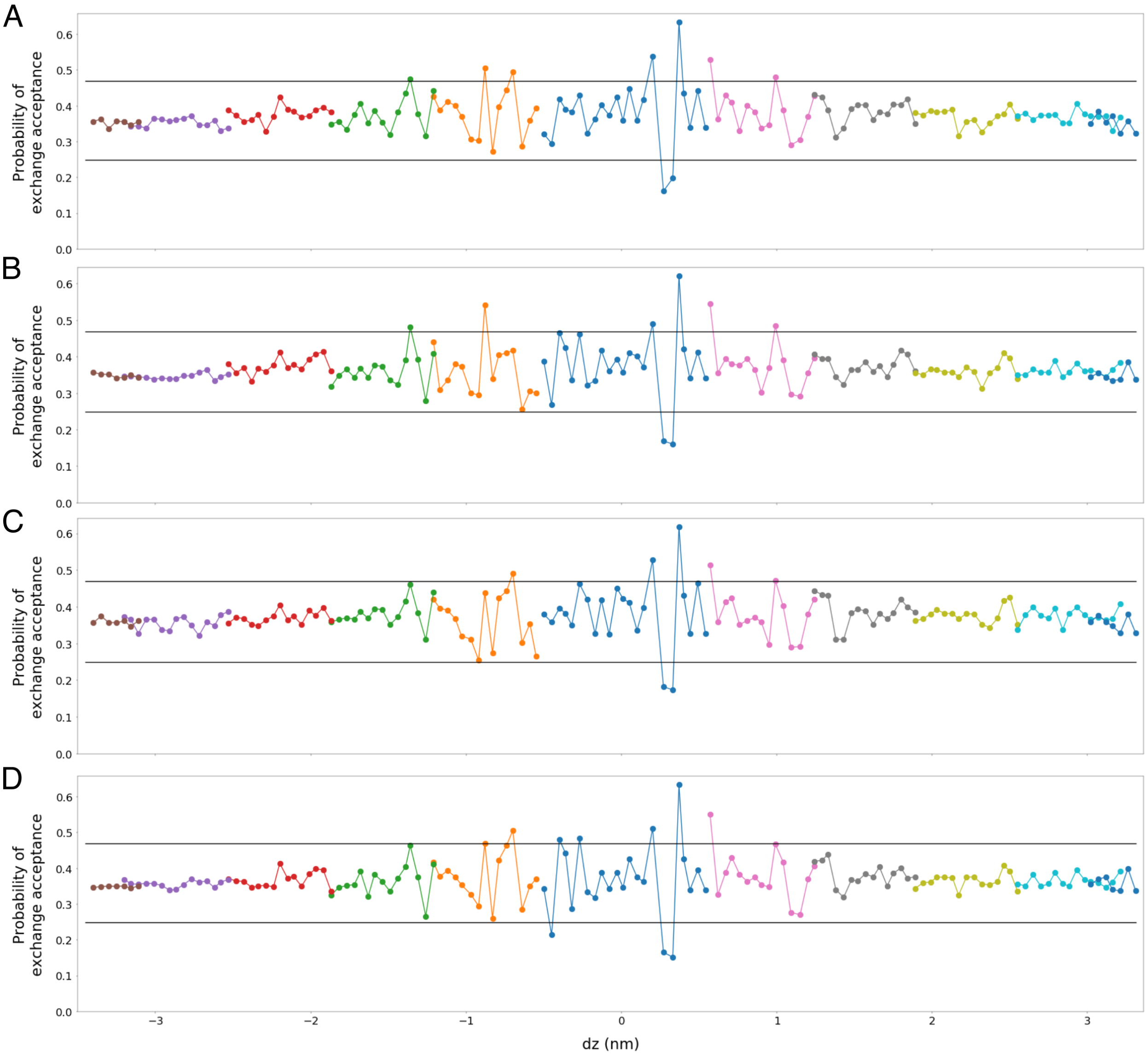


Figure S6: Average exchange probabilities between umbrella windows after 200 ns of production simulation for two forward replicates (A–B), and two reverse transition (C–D) in orientation 2. For more details, see legend of Fig. S5
